# PEG Adjustment Enables Genome Transplantation Using *Mycoplasma Mycoides* Recipient

**DOI:** 10.1101/2024.09.27.615432

**Authors:** Bogumil J. Karas, Nicolette G. Moreau, J Craig Venter, Hamilton O Smith, John I. Glass

## Abstract

Pioneering advances in synthetic biology, initiated at the J. Craig Venter Institute, have enabled the creation of the first cell driven by synthetic genome and opened a new era of creating designer microbes. This technology offers far greater potential for genetic modification than traditional genome editing techniques and holds promise for the routine creation of synthetic cells as DNA synthesis costs decrease and genetic knowledge expands. Two essential technologies underpin this achievement: the ability to clone entire genomes in host organisms, such as *Saccharomyces cerevisiae*, and the successful transplant of these genomes into recipient cells to generate living organisms. In all previous work the recipient cell in genome transplantation experiments has been *Mycoplasma capricolum*. In this study, we explored the potential of using *Mycoplasma mycoides* strains as recipient cells for genome transplantation. By increasing polyethylene glycol (PEG) concentration from 5% to 10%, we successfully transplanted various *M. mycoides* strains using several *M. mycoides* recipient cells. Additionally, we demonstrated the ability to transplant *M. capricolum* genomes into *M. mycoides* recipient cells; although with lower efficiency compared to *M. mycoides* strains. These findings provide a modified transplantation protocol and likely get us closer to expanding genome transplantation to other bacterial species.

## Introduction

Pioneering technology developed at the J. Craig Venter Institute led to the creation of the first cell driven by synthetic genome in 2010 ^1^. This historic achievement marked the beginning of an era of creating designer life. This technology holds tremendous promise, enabling virtually limitless genetic modifications that are impossible with genome editing technologies like CRISPR/Cas9 ^2^. As DNA synthesis costs drop and genetic knowledge advances, creating synthetic cells could become a routine process, accessible even to undergraduate students conducting lab experiments. Such access could accelerate the development of industrial microbes with specialized functions or strains designed for studying fundamental cellular processes.

Two key technologies had to be developed to enable the creation of synthetic cells: 1) cloning whole genomes in host organisms and 2) transplanting or “booting up” these designer genomes to create living cells. The best host for genome cloning is baker’s yeast (*Saccharomyces cerevisiae*). Benders and colleagues described three approaches to assembling whole genomes ^3^, which can be done from synthetic fragments or a combination of synthetic fragments and cloned genome fragments. Lartigue et al., in an elegant experiment, demonstrated successful transplantation of the entire genome of *Mycoplasma mycoides* into *Mycoplasma capricolum* ^4,5^. Some of the key features of this pair of mycoplasmas are that they had small genomes (∼1 Mbp) that are phylogenetically very closely related, compatible restriction-modification systems and a lack of cell wall ^4^.

Since then, the most prominent example of this technology has been the creation of cells with a minimized genome ^1^. The transplantation process is one of the biggest challenges in expanding this technology to other organisms. So far, all transplantations have been performed using *M. capricolum* recipient cells, with the following donor strains: *Mycoplasma leachii* PG50, *Mycoplasma putrefaciens* strain 156, *Mycoplasma forum* L1, *M. capricolum* subsp. *capricolum* CK ^6^, *M. capricolum* subsp. *Capripneumoniae* ^7^ and the fast-growing *Mycoplasma feriruminatoris* ^8^. At the same time, whole genomes from *Acholeplasma laidlawii, Prochlorococcus marinus*, and *Haemophilus influenzae* have been cloned in yeast ^9–11^ but limited efforts to transplant those genomes failed. A primary challenge in bacterial genome transplantation lies in successfully transferring the donor genome into the recipient cell. This process is still poorly understood and requires further investigation before being widely adopted. Figure 1 below summarizes the current transplantation protocol and suggests future tests to overcome the transplantation barriers.

**Figure 1.**
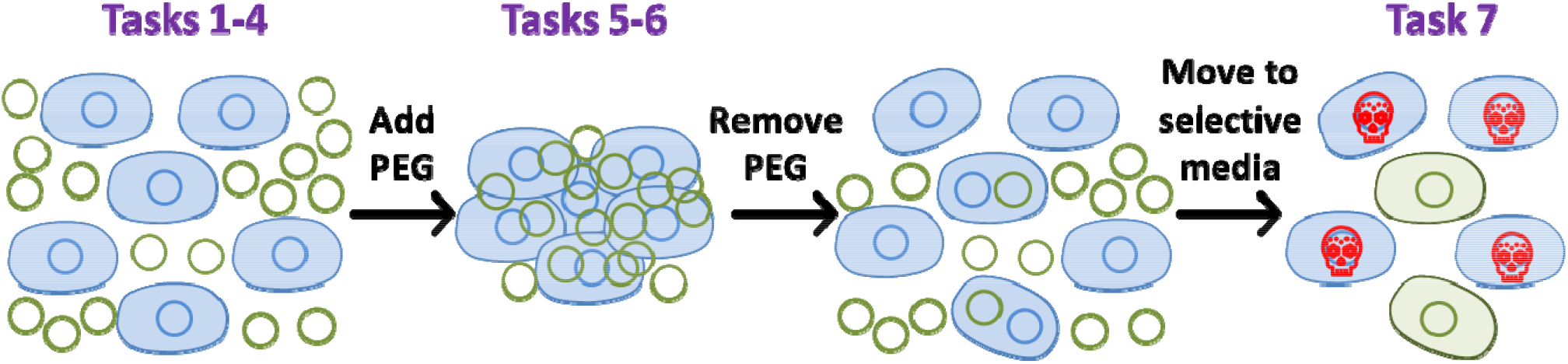
Proposed mechanisms of genome transplantation. In the original protocol ^4^, the donor DNA (green circles) is mixed with recipient cells (blue) in a 0.1 M CaCl_2_ solution, followed by incubation in 5% polyethylene glycol (PEG). This mixture facilitates DNA/recipient cell aggregation, likely leading to cell fusion. Some donor DNA can be trapped inside the recipient cells during fusion. Afterward, the PEG solution is removed, allowing cells to recover, and media containing an antibiotic specific to the donor genome is introduced. Only successfully transplanted genomes or potential hybrid genomes survive this selection process (the red skull indicates dead cells). Recommended/necessary tasks for adapting this process to other species: 1) Developing a recipient strain free of restriction and recombination systems; 2) Modifying the membrane/cell wall of recipient cells through growth conditions (such as temperature), chemical treatment (e.g., removal of the cell wall), or genetic modifications; 3) Development of improved method for preparing intact donor genomic DNA; 4) Testing various growth temperatures when preparing recipient cells; 5) Testing various concentrations and molecular weights of PEG; 6) Testing various incubation time in the PEG solution; 7) Evaluating different antibiotic concentrations. Note: Figure adopted from ^12^.

In this work, we investigated whether *M. mycoides* strains could be used as a recipient cell. We demonstrated that by increasing PEG concentration from 5% to 10%, we were able to successfully transplant multiple *M. mycoides* strains in a few different *M. mycoides* recipient cells. Additionally, we showed that it is possible to transplant *M. capricolum*, although not as efficiently as when *M. mycoides* donor strains are used. We provided a modified transplantation protocol, representing an important step toward expanding genome transplantation to other species.

## Results and Discussion

To develop the “reverse” transplantation protocol using *M. mycoides* as the recipient, we initially tested conditions published for using *M. capricolum* as the recipient ^4^. Our donor was a pool of wild-type *M. mycoides* carrying a randomly inserted puromycin-resistant gene, and the recipient was the synthetic version of *M. mycoides*: JCVI-syn1.0, which carries a tetracycline-resistant gene. After numerous trials, we were unsuccessful. One of the critical steps in the published protocol involves incubating the recipient cells and donor DNA in 5% PEG. We next investigated if changing PEG concentration will have any effect on genome transplantation in the reverse direction (refers to using *M. mycoides* as recipient instead of *M. capricolum*). We tested 5% and 10% PEG 8000 concentrations, along with varying incubation times from 90 minutes up to 210 minutes. For all incubation times we obtained colonies only with 10% PEG (Figure 2A).

**Figure 2.**
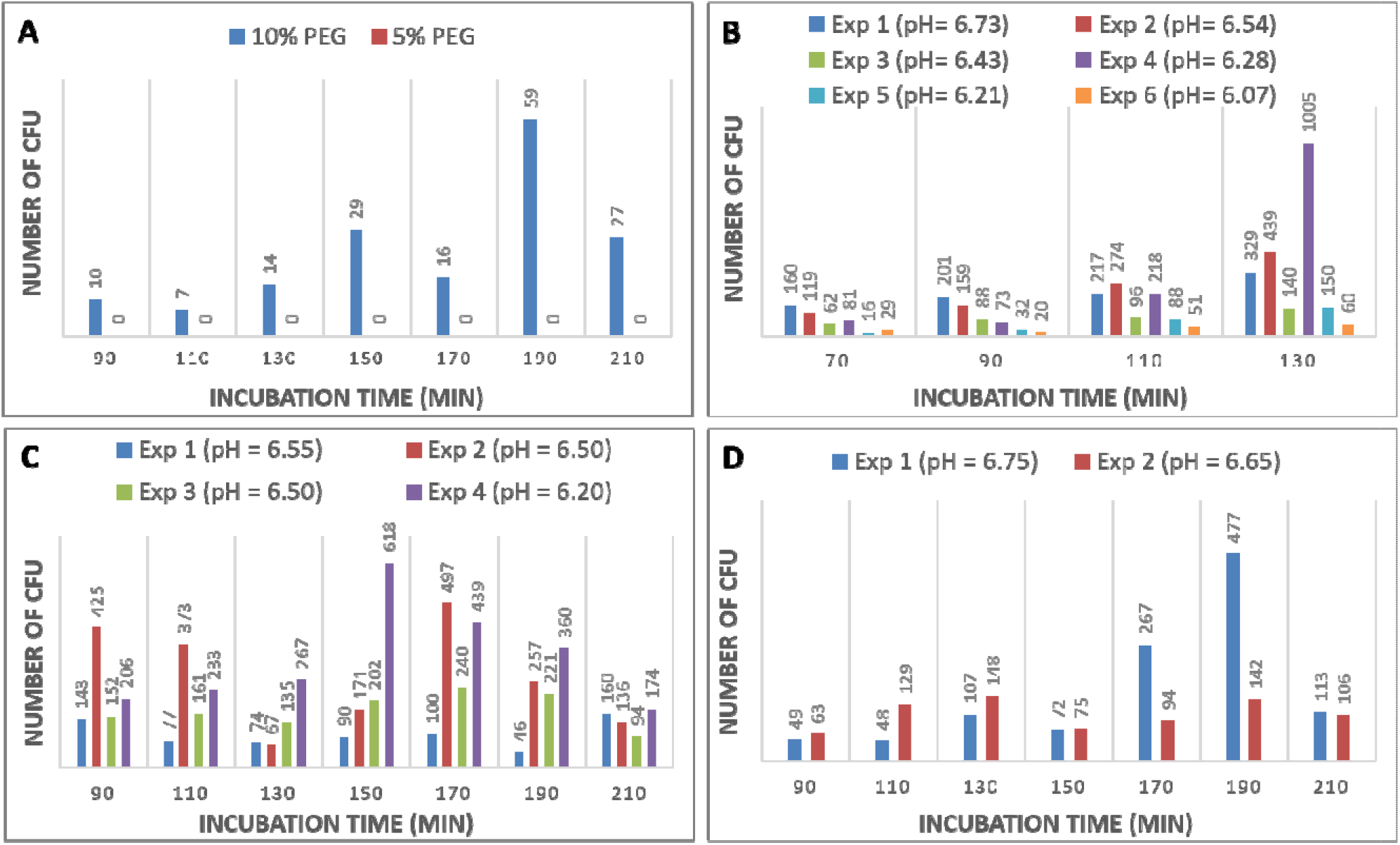
Reverse transplantation experiments. For all experiments, CFU indicates colony-forming units. **A)** Testing 5% vs 10% PEG, as well as incubation time. Donor: Wild type *M. mycoides* carrying puromycin resistance gene, Recipient: JCVI-Syn1.0 carrying tetracycline resistance gene. Note: Recipient cells had pH = 6.58 at harvest. **B)** First set of experiments testing recipient cells culture pH at harvest, as well as incubation time. Donor: Wild type *M. mycoides* carrying puromycin resistance gene, Recipient: partially reduced M. myoides strain RGD1-2 carrying tetracycline resistance gene. **C)** Second set of experiments testing recipient cells culture pH at harvest, as well as incubation time. Donors and recipients are the same as in B. **D)** Third set of experiments testing recipient cells culture pH at harvest, as well as incubation time. Donor: JCVI-Syn1.0 carrying puromycin resistance gene, Recipient: JCVI-Syn1.0 restriction minus, carrying tetracycline resistance gene.

Following this successful result, we tested three different recipient cells with partially minimized genomes RGD1-2, RGD1-3, and RGD1-6, where each strain has one-eighth of the genome minimized ^1^. These strains were selected as they produced the best results in related experiments, where genomes are transferred via cell fusion/engulfment from bacteria to yeast ^13^. The best recipient strain was RGD1-2 (preliminary data not shown). We then proceeded with this strain, testing different culture pH levels and incubation times in PEG, ranging from 70 to 130 minutes (Figure 2B). For all six experiments, the best data was obtained only for 130 minutes of incubation (Figure 2B). Consequently, we conducted another set of experiments with incubation times extended from 90 to 210 minutes (Figure 2C). The optimal incubation times were 170 minutes for two experiments, 150 minutes for one, and 210 minutes for another (Figure 2C). We then replicated the same conditions with two additional experiments using the JCVI-Syn1.0 restriction minus recipient (2D). For both experiments, the best results were achieved at 190 minutes of incubation, showing ∼ 10x or ∼2x improvement compared to the 90-minute incubation (2D). The detailed reverse transplantation protocol is listed in Figure 3.

**Figure 3.**
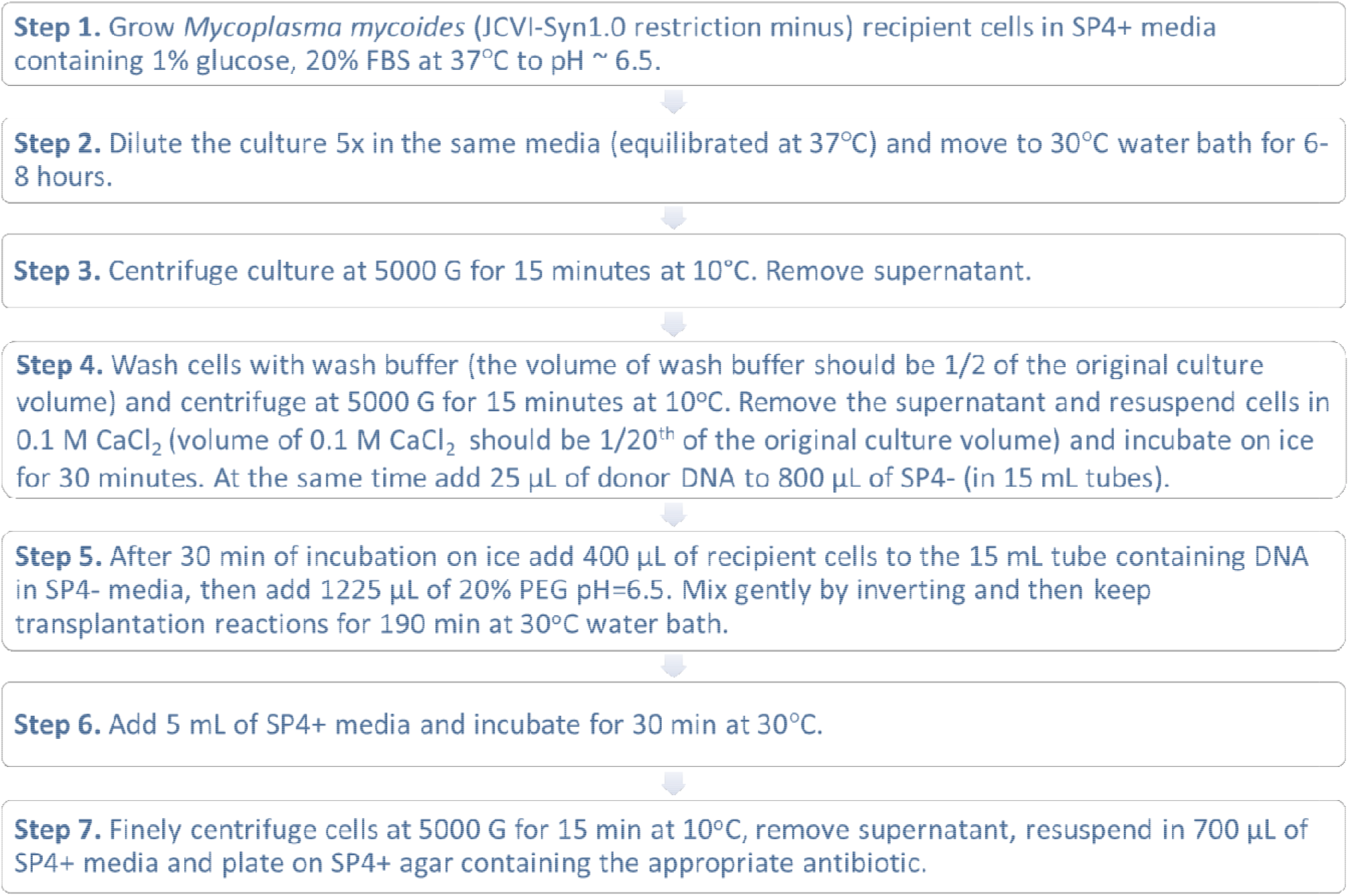
Reverse genome transplantation protocol for using *M. mycoides* recipient cells. Note that it works for transplanting compatible M. mycides and *M. capricolum* donor strains (see below).

We then attempted to transplant the *M. capricolum* carrying puromycin gene into JCVI-Syn1.0 restriction-modification minus strains using the new protocol. Multiple experiments were conducted, but only a few produced *M. capricolum* colonies (Figure 4A) while obtaining a high number of colonies for JCVI-Syn1.0 donor using the same preparation of recipient cells. Since the transplantation protocol for using *M. capricolum* as recipient cells requires 5% PEG, we hypothesize that 10% PEG could have a negative effect on *M. capricolum* cells. To test this, we incubated both donor strains from Figure 4A for varying amounts of time in 5% and 10% PEG, as shown in Figures 4B and 4C. Interestingly, for JCVI-Syn1.0 no differences were observed, but there was close to a 50% drop in CFU for *M. capricolum* after 90 minutes, and this trend remained consistent across all times tested. Additional experiments will need to be performed to optimize the transplantation of *M. capricolum* in JCVI-Syn1.0 restriction-minus strains.

**Figure 4.**
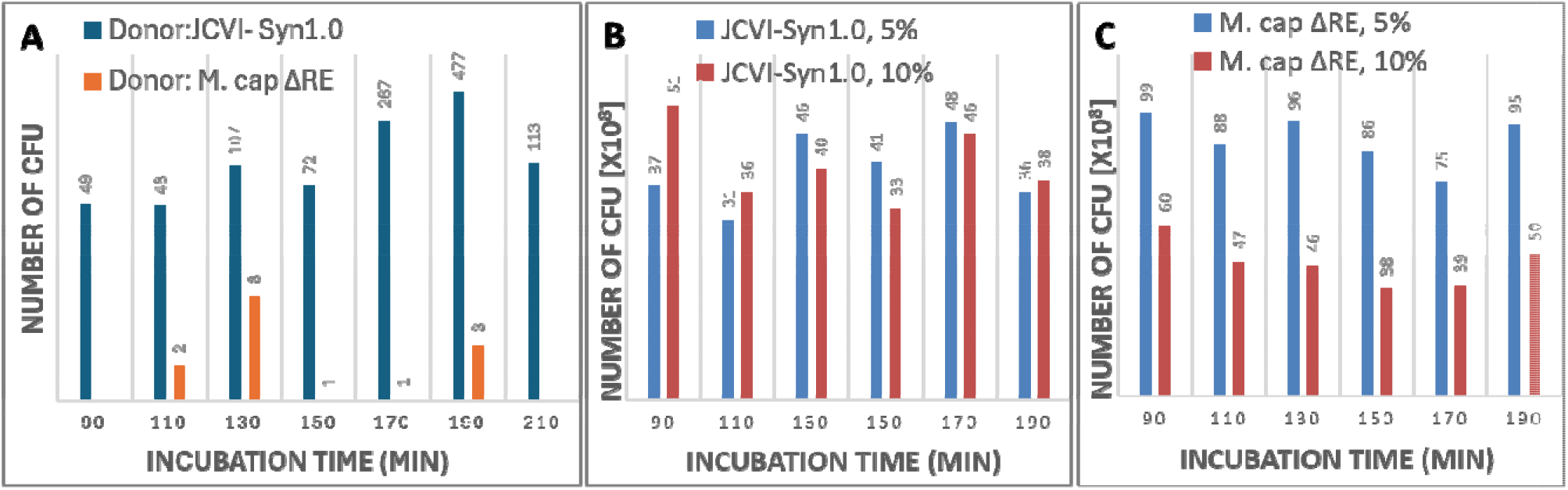
Transplantation of *M. capricolum*. **A)** Transplantation of *M. capricolum* restriction minus strain carrying puromycin resistant gene using JCVI-syn1.0 restriction minus recipient cells. As a control JCVI-Syn1.0 donor was used. Note: the data for JCVI-Syn1.0 is also shown in Figure 2D. **B)** Survival of JCVI-Syn1.0 cells in 5 and 10% PEG. C) Survival of JCVI-Syn1.0 cells in 5 and 10% PEG.

## Final conclusions

In conclusion, this study demonstrates a significant advancement in genome transplantation technology by successfully modifying the established protocol for *Mycoplasma mycoides* recipient cells. By increasing the concentration of polyethylene glycol (PEG) from 5% to 10%, we achieved successful transplantation of multiple *M. mycoides* strains and demonstrated limited success with *M. capricolum*. This modified protocol provides a foundation for future exploration of genome transplantation in other bacterial species and opens the door for further optimization of genome transfer techniques.

## METHODS

### Strains used and cultured conditions

All *Mycoplasma* strains ^1,5,14^ were cultured in SP-4 ^15^.

### Creation of transposon insertion strains of M mycoides and *M. capricolum*

As described in ^1^

### Transplantation protocol

As described in ^4^ with the modifications as described in Figure 3 in this manuscript

## Supporting Information

## ACKNOWLEDGEMENT

This work was supported by Synthetic Genomics, Inc. and the US National Science Foundation grants MCB-1840301 and MCB-2218507. In addition, research by B.J.K is supported by the Natural Sciences and Engineering Research Council of Canada (RGPIN-2018-06172).

## Author Contributions

B.J.K., N.G.M., performed the experiments.; B.J.K., N.G.M., J.C.V., H.O.S., J.I.G., designed experiments and interpreted results; B.J.K. wrote the paper; B.J.K., N.G.M., J.I.G., edited the final version.

## References

(1) Hutchison, C. A., III; Chuang, R.-Y.; Noskov, V. N.; Assad-Garcia, N.; Deerinck, T. J.; Ellisman, M. H.; Gill, J.; Kannan, K.; Karas, B. J.; Ma, L.; Pelletier, J. F.; Qi, Z.-Q.; Richter, R. A.; Strychalski, E. A.; Sun, L.; Suzuki, Y.; Tsvetanova, B.; Wise, K. S.; Smith, H. O.; Glass, J. I.; Merryman, C.; Gibson, D. G.; Venter, J. C. Design and Synthesis of a Minimal Bacterial Genome. Science 2016, 351 (6280), aad6253–aad6253.

(2) Jinek, M.; Chylinski, K.; Fonfara, I.; Hauer, M.; Doudna, J. A.; Charpentier, E. A Programmable Dual-RNA–Guided DNA Endonuclease in Adaptive Bacterial Immunity. Science 2012, 337 (6096), 816–821.

(3) Benders, G. A.; Noskov, V. N.; Denisova, E. A.; Lartigue, C.; Gibson, D. G.; Assad-Garcia, N.; Chuang, R.-Y.; Carrera, W.; Moodie, M.; Algire, M. A.; Phan, Q.; Alperovich, N.; Vashee, S.; Merryman, C.; Venter, J. C.; Smith, H. O.; Glass, J. I.; Hutchison, C. A., 3rd. Cloning Whole Bacterial Genomes in Yeast. Nucleic Acids Res. 2010, 38 (8), 2558–2569.

(4) Lartigue, C.; Glass, J. I.; Alperovich, N.; Pieper, R.; Parmar, P. P.; Hutchison, C. A., 3rd; Smith, H. O.; Venter, J. C. Genome Transplantation in Bacteria: Changing One Species to Another. Science 2007, 317 (5838), 632–638.

(5) Lartigue, C.; Vashee, S.; Algire, M. A.; Chuang, R.-Y.; Benders, G. A.; Ma, L.; Noskov, V. N.; Denisova, E. A.; Gibson, D. G.; Assad-Garcia, N.; Alperovich, N.; Thomas, D. W.; Merryman, C.; Hutchison, C. A., 3rd; Smith, H. O.; Venter, J. C.; Glass, J. I. Creating Bacterial Strains from Genomes That Have Been Cloned and Engineered in Yeast. Science 2009, 325 (5948), 1693– 1696.

(6) Labroussaa, F.; Lebaudy, A.; Baby, V.; Gourgues, G.; Matteau, D.; Vashee, S.; Sirand-Pugnet, P.; Rodrigue, S.; Lartigue, C. Impact of Donor-Recipient Phylogenetic Distance on Bacterial Genome Transplantation. Nucleic Acids Res. 2016, 44 (17), 8501–8511.

(7) A Toolbox for Manipulating the Genome of the Major Goat Pathogen, Mycoplasma Capricolum Subsp. Capripneumoniae Géraldine Gourgues, Lucía Manso-Silván; Catherine Chamberland, Pascal Sirand-Pugnet, François Thiaucourt, Alain.

(8) Talenton, V.; Baby, V.; Gourgues, G.; Mouden, C.; Claverol, S.; Vashee, S.; Blanchard, A.; Labroussaa, F.; Jores, J.; Arfi, Y.; Sirand-Pugnet, P.; Lartigue, C. Genome Engineering of the Fast-Growing Mycoplasma Feriruminatoris toward a Live Vaccine Chassis. ACS Synth. Biol. 2022, 11 (5), 1919–1930.

(9) Karas, B. J.; Tagwerker, C.; Yonemoto, I. T.; Hutchison, C. A., 3rd; Smith, H. O. Cloning the Acholeplasma Laidlawii PG-8A Genome in Saccharomyces Cerevisiae as a Yeast Centromeric Plasmid. ACS Synth. Biol. 2012, 1 (1), 22–28.

(10) Karas, B. J.; Jablanovic, J.; Sun, L.; Ma, L.; Goldgof, G. M.; Stam, J.; Ramon, A.; Manary, M. J.; Winzeler, E. A.; Venter, J. C.; Weyman, P. D.; Gibson, D. G.; Glass, J. I.; Hutchison, C. A., 3rd; Smith, H. O.; Suzuki, Y. Direct Transfer of Whole Genomes from Bacteria to Yeast. Nat. Methods 2013, 10 (5), 410–412.

(11) Tagwerker, C.; Dupont, C. L.; Karas, B. J.; Ma, L.; Chuang, R.-Y.; Benders, G. A.; Ramon, A.; Novotny, M.; Montague, M. G.; Venepally, P.; Brami, D.; Schwartz, A.; Andrews-Pfannkoch, C.; Gibson, D. G.; Glass, J. I.; Smith, H. O.; Venter, J. C.; Hutchison, C. A., 3rd. Sequence Analysis of a Complete 1.66 Mb Prochlorococcus Marinus MED4 Genome Cloned in Yeast. Nucleic Acids Res. 2012, 40 (20), 10375–10383.

(12) Karas, B. J.; Suzuki, Y.; Weyman, P. D. Strategies for Cloning and Manipulating Natural and Synthetic Chromosomes. Chromosome Res. 2015, 23 (1), 57–68.

(13) Karas, B. J.; Moreau, N. G.; Deerinck, T. J.; Gibson, D. G.; Venter, J. C.; Smith, H. O.; Glass, J. I. Direct Transfer of a Mycoplasma Mycoides Genome to Yeast Is Enhanced by Removal of the Mycoides Glycerol Uptake Factor Gene GlpF. ACS Synth. Biol. 2019, 8 (2), 239–244.

(14) Gibson, D. G.; Glass, J. I.; Lartigue, C.; Noskov, V. N.; Chuang, R.-Y.; Algire, M. A.; Benders, G. A.; Montague, M. G.; Ma, L.; Moodie, M. M.; Merryman, C.; Vashee, S.; Krishnakumar, R.; Assad-Garcia, N.; Andrews-Pfannkoch, C.; Denisova, E. A.; Young, L.; Qi, Z.-Q.; Segall-Shapiro, T. H.; Calvey, C. H.; Parmar, P. P.; Hutchison, C. A., 3rd; Smith, H. O.; Venter, J. C. Creation of a Bacterial Cell Controlled by a Chemically Synthesized Genome. Science 2010, 329 (5987), 52– 56.

(15) Tully, J. G.; Rose, D. L.; Whitcomb, R. F.; Wenzel, R. P. Enhanced Isolation of Mycoplasma Pneumoniae from Throat Washings with a Newly-Modified Culture Medium. The Journal of infectious diseases 1979, 139 (4), 478–482.

